# Apoptotic cells promote circulating tumor cell survival and metastasis

**DOI:** 10.1101/2024.05.21.595217

**Authors:** Cassidy E. Hagan, Annelise G. Snyder, Mark Headley, Andrew Oberst

## Abstract

During tumor progression and especially following cytotoxic therapy, cell death of both tumor and stromal cells is widespread. Despite clinical observations that high levels of apoptotic cells correlate with poorer patient outcomes, the physiological effects of dying cells on tumor progression remain incompletely understood. Here, we report that circulating apoptotic cells robustly enhance tumor cell metastasis to the lungs. Using intravenous metastasis models, we observed that the presence of apoptotic cells, but not cells dying by other mechanisms, supports circulating tumor cell (CTC) survival following arrest in the lung vasculature. Apoptotic cells promote CTC survival by recruiting platelets to the forming metastatic niche. Apoptotic cells externalize the phospholipid phosphatidylserine to the outer leaflet of the plasma membrane, which we found increased the activity of the coagulation initiator Tissue Factor, thereby triggering the formation of platelet clots that protect proximal CTCs. Inhibiting the ability of apoptotic cells to induce coagulation by knocking out Tissue Factor, blocking phosphatidylserine, or administering the anticoagulant heparin abrogated the pro-metastatic effect of apoptotic cells. This work demonstrates a previously unappreciated role for apoptotic cells in facilitating metastasis by establishing CTC-supportive emboli, and suggests points of intervention that may reduce the pro-metastatic effect of apoptotic cells.

**GRAPHICAL ABSTRACT:** 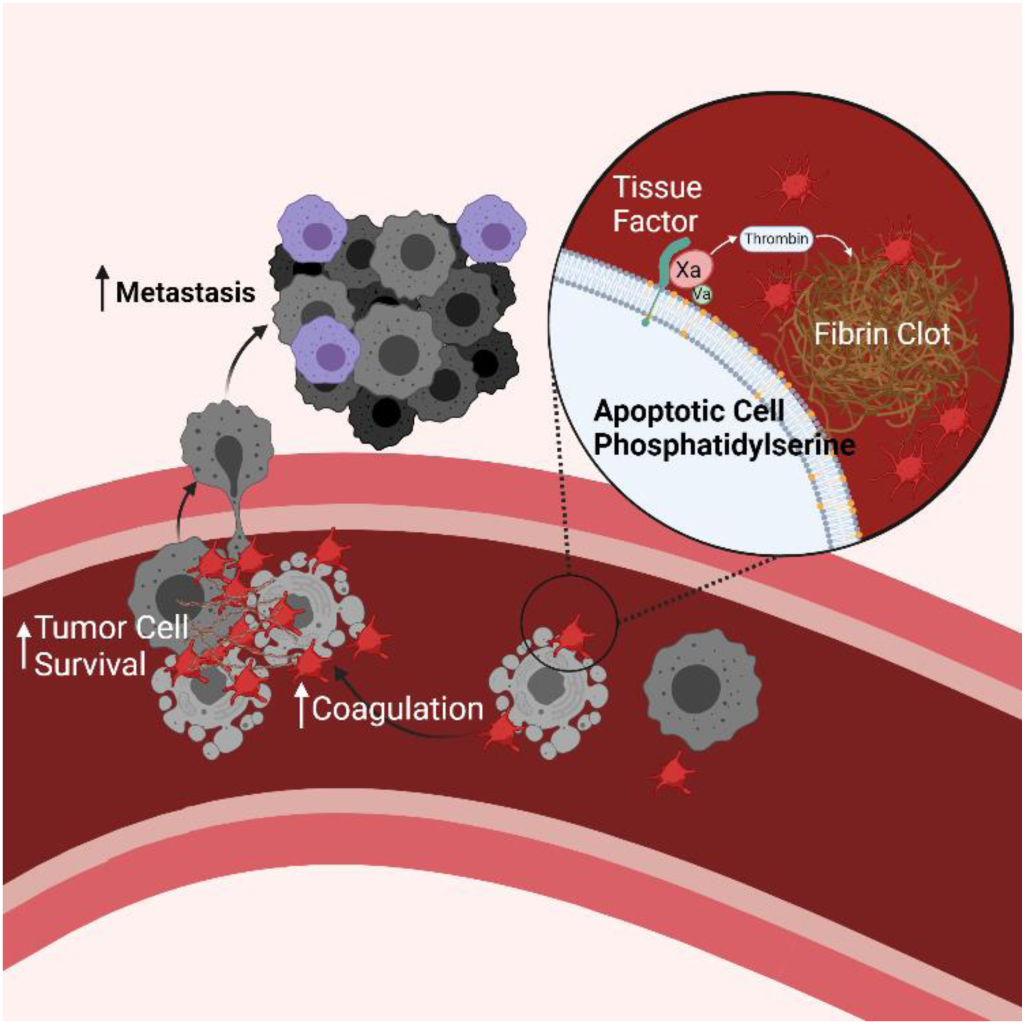

## INTRODUCTION

Metastatic disease, caused by the dissemination of tumor cells from a primary tumor to distant tissues, is the major cause of mortality in cancer patients (1). Efforts to understand the process of metastasis have identified key features of circulating tumor cells (CTCs) that successfully seed distant tissues and defined aspects of the microenvironment required to support CTCs throughout the metastatic cascade (2). CTCs must create a supportive niche to allow their survival in the hostile circulatory environment and upon arrest at distant tissue sites. A key feature of this process is the activation of coagulation by CTCs; this leads to the coating of CTCs by activated platelets, which provide survival signals and protect CTCs from shear stress and natural killer (NK) cell recognition (3–9). Platelet activation is required for the subsequent recruitment of neutrophils and monocytes (10,11) and increases endothelial permeability to support extravasation (12). Further underscoring the importance of cell-cell interactions, CTC clusters have significantly enhanced metastatic potential when compared to individual CTCs (13–16).

Overall, the metastatic process is extremely inefficient. Experiments tracking the fate of CTCs have found that <1% of cells survive the stresses associated with vascular dissemination (17,18). Recognition by immune cells as well as shear stress leads to the demise of a majority of CTCs before they can arrest, extravasate and form a metastatic tumor. While most research has focused on characteristics that allow rare CTCs to successfully metastasize, very little is understood about the effects of the greater portion of tumor cells that die during the process of metastasis.

Studies examining CTCs have found that a large proportion show molecular features of apoptosis, and in patients with breast cancer, the presence of apoptotic CTCs is correlated with progression to metastatic disease (19–23). Furthermore, apoptosis within primary tumors has been shown to promote tumor growth via multiple mechanisms, including promoting selection for aggressive cellular clones (24), releasing proliferative signals (25), and recruiting and polarizing macrophage populations (26,27). However, experimental evidence evaluating the effect of circulating apoptotic cells on metastasis is lacking.

Apoptosis is executed following the activation of caspases, a family of protease enzymes whose targets define the morphological changes associated with apoptotic cell death. Caspase 8 is activated following extrinsic stimuli, while caspase 9 is activated downstream of mitochondrial permeabilization during intrinsic apoptosis. Each pathway converges on the activation of executioner caspases 3 and 7 which are responsible for the highly ordered demise of the cell. Numerous caspase targets exist, resulting in canonical features of apoptosis such as cell shrinkage, membrane blebbing, externalization of phosphatidylserine from the inner to the outer leaflet of the plasma membrane, DNA degradation, and chromatin condensation. Apoptosis is considered an anti-inflammatory form of programmed cell death, inducing programs of tolerance and wound healing, consistent with its putative role in promoting tumor growth (28–30).

Here, we show that apoptotic cells in circulation potently enhance metastasis by promoting the survival of CTCs. This effect requires the physical association of apoptotic cells with CTCs at sites of metastatic colonization, and is driven by the promotion of coagulation and platelet aggregation by apoptotic cells. The procoagulant activity of apoptotic cells is dictated by Tissue Factor (TF) activation, which occurs upon phosphatidylserine externalization on the apoptotic cell surface. The ability of apoptotic cells to promote metastasis correlates with the colocalization of tumor cells and apoptotic cells within platelet clots, establishing apoptotic cells as an important contributor to the early metastatic microenvironment supporting tumor cell survival in circulation.

## RESULTS

### Circulating apoptotic cells increase lung metastasis in I.V. metastasis models

Given the clinical correlation between apoptotic CTCs and progression to metastasis in patients with breast cancer (19,20), we sought to design a system in which we could directly assess the impact of apoptotic cells during hematogenous dissemination of breast cancer. To do this, Met-1 cells, a mammary tumor cell line derived from MMTV-PyMT transgenic mice (31), were injected intravenously (I.V.) into syngeneic (FVBN/J) immunocompetent mice at a 1:1 ratio with apoptotic cells. Apoptotic cells were prepared from cells lines expressing caspase-8 or caspase-9 fused to activatable FKBP^F36V^ dimerization domains (hereafter acCasp8, acCasp9). Incubating these cells with the nontoxic ligand B/B, a synthetic bivalent homolog of rapamycin, enforces caspase dimerization and activation. This system, which we and others have used in the past, gives us precise control over the activation of apoptosis (32–35). After a short pulse incubation with B/B *ex vivo* cells are committed to undergo apoptosis and are immediately processed for I.V. injections such that apoptotic cells undergo the apoptotic cascade *in vivo.* Inducing apoptosis through acCasp9 resulted in rapid externalization of phosphatidylserine (PS) while acCasp8 activation or UV irradiation resulted in slower, more heterogeneous apoptotic cell death (**Fig. 1A)**. In all cases, induction of apoptosis led to 100% cell death as measured by membrane permeability **(Sup. 1)**. Fourteen days following I.V. injection of Met-1 cells alone or Met-1 cells mixed with dying cells, mice were euthanized and the number of metastatic nodules on the lung surface was quantified **(Fig. 1A)**.

**Figure 1.**
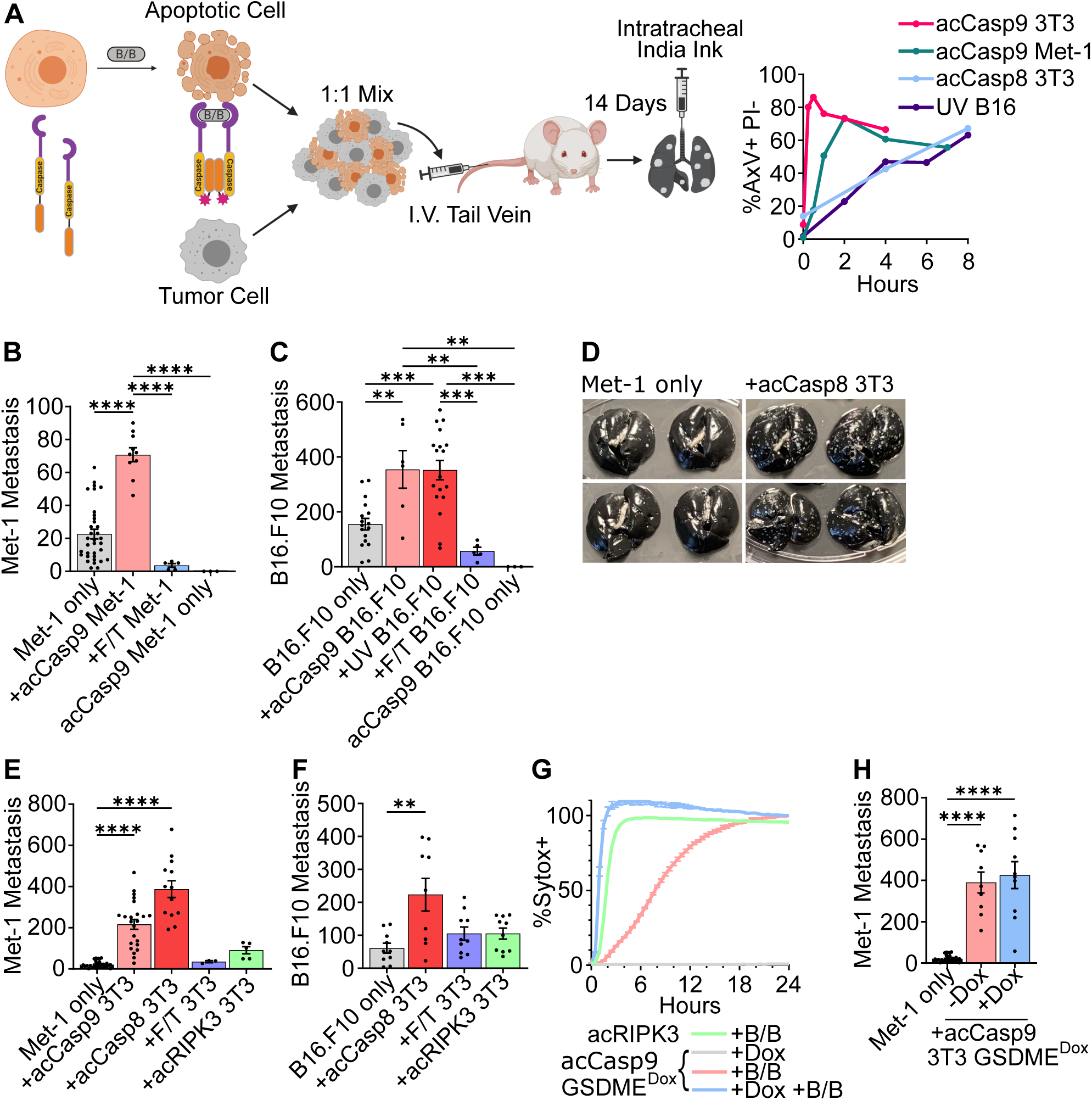
Circulating apoptotic cells increase lung metastasis in I.V. metastasis models. acCasp8, acCasp9, or acRIPK3 expressing cells were activated by treatment with B/B for 15 minutes, UV irradiated, or killed by Freeze/Thaw lysis (F/T). PS externalization and membrane integrity on cells after B/B or UV exposure was measured with Annexin V (AxV) and propidium iodide (PI) by flow cytometry (A). Dying cells were mixed 1:1 with tumor cells and injected I.V., 14 days later surface lung metastasis was quantified (A-F, H). GSDME expression in acCasp9 3T3 GSDME^Dox^ cells was induced by doxycycline treatment (+Dox), followed by treatment with B/B to activate acCasp9 (G-H). Membrane permeability was measured by Sytox Green uptake (G). Dots are biological replicates, Error bars represent SEM, statistical testing is Ordinary one-way ANOVA with Tu key’s multiple comparisons test.

We observed that coinjection of apoptotic Met-1 cells with viable Met-1 cells increased the metastatic burden by approximately three-fold, while necrotic Met-1 cells killed via freeze/thaw (F/T) lysis had no impact on metastasis **(Fig. 1B)**. We also examined a model of melanoma metastasis, B16.F10 cells injected I.V. into B6/J mice, and found a similar increase in metastasis with coinjection of apoptotic but not necrotic tumor cells **(Fig. 1C)**. Induction of apoptosis in B16.F10 cells through both UV irradiation and acCasp9 activation had similar effects. We confirmed that apoptotic cells themselves could not form metastatic nodules; injecting only apoptotic cells resulted in no detectable tumor foci in the lung (**Fig. 1B- C**).

To examine whether this was a general feature of apoptotic cells regardless of cell type, we tested the effect of coinjecting apoptotic NIH-3T3 fibroblasts (3T3) in Met-1 and B16.F10 metastasis models. Strikingly, apoptotic 3T3s enhanced Met-1 metastasis by approximately 20-fold **(Fig. 1D-E).** Apoptotic 3T3s had a similar effect on B16.F10 metastasis as apoptotic B16.F10 cells **(Fig. 1F)**.

We confirmed that the allogenicity of 3T3s was not a relevant variable by observing that autologous mouse embryonic fibroblasts (MEF) stimulated to undergo apoptosis with UV irradiation also enhanced Met-1 and B16.F10 metastasis to a similar degree as apoptotic 3T3s **(Sup. 2A-B)**. Our finding that allogeneic and autologous apoptotic fibroblasts as well as apoptotic tumor cells enhance metastasis suggests that the promotion of metastasis is a general feature of apoptotic cells, regardless of their origin.

As observed with necrotic Met-1 and B16.F10, necrotic 3T3 fibroblasts had no effect on metastasis in either model **(Fig. 1E-F)**. Inducing necroptosis, a form of programmed lytic cell death, through acRIPK3 oligomerization in 3T3 cells (32,35) also had no impact on metastasis. The caspase-3 cleavable pore-forming protein Gasdermin E (GSDME) is responsible for secondary necrosis following apoptosis (36). Doxycycline (Dox) controlled overexpression of GSDME, caused cells to progress to secondary necrosis rapidly upon caspase activation, comparable to the rate of lysis after acRIPK3 oligomerization **(Fig. 1G)**. However, GSDME expression did not impact the ability of acCasp9 cells to enhance metastasis. Together this suggests that caspase-independent lytic cell death does not promote metastasis, however, cell lysis following caspase activation does not terminate the pro-metastatic effect of apoptotic cells **(Fig. 1H)**.

### Apoptotic cells provide a survival advantage to circulating tumor cells

We next sought to address where, along the metastatic cascade, apoptotic cells exerted their prometastatic effects. We first examined whether apoptosis might act by suppressing adaptive immune surveillance of metastatic cells by examining their effect in *Rag2^-/-^* mice, which lack T and B cells. Apoptotic cells retained their ability to promote metastasis in *Rag2^-/-^* mice, indicating that the adaptive immune compartment was not a primary driver of the effects of apoptotic cells **(Sup. 3A)**. Next, to examine the contribution of NK cells which play an important role in restricting CTC survival (37,38), we administered an NK depletion antibody (anti-NK1.1) 72 and 24 hours prior to I.V. metastasis challenge. Although depleting NK cells resulted in substantially higher metastatic burden, as previously reported (39), apoptotic cells still provided a metastatic advantage, suggesting that apoptotic cells are acting via a mechanism other than protecting tumor cells from NK cell killing. **(Sup. 3B-C)**

We next sought to assess when during the dynamic process of tumor cell arrest, survival and extravasation apoptotic cells exert their pro-metastatic effect. To quantify tumor cell seeding and subsequent survival we used qPCR to measure tumor-specific genomic DNA (PyMT transgene) in the lung at timepoints from 5 minutes to 96 hours post I.V. injection. Maximum tumor cell quantity in the lung was observed as early as 5 minutes following I.V. injection, indicating rapid arrest of tumor cells in the lung vasculature. This initial seeding was not affected by the co-administration of apoptotic cells. The relative quantity of tumor cells declined exponentially over the next 24 hours as previously reported (18) **(Fig. 2A-B).** We concurrently quantified apoptotic cell genomic DNA in the lung and as expected apoptotic cells showed a similar logarithmic decline in abundance over 24 hours, concomitant with their uptake by CD45+ phagocytes **(Sup. 4A-C)**. We found that tumor cell survival (as measured by persistence of tumor genomic material over the first 24 hours) was increased in the presence of apoptotic cells. Coinjection of Met-1 tumor cells with apoptotic 3T3 cells resulted in an increase in PyMT gDNA (Met-1 cells) of approximately 4-fold at 6 hours post-injection and 50-fold at 24 hours post-injection **(Fig. 2A)**. Met-1 cells expressing ZsGreen to distinguish them from coinjected apoptotic acCasp9 Met-1 cells also showed increased gDNA persistence in the lungs at 6 and 24 hours post-injection relative to Met-1 cells injected alone **(Fig. 2B)**. We observed similar trends when quantifying tumor cells in the lung by flow cytometry and by fluorescent imaging: apoptotic cells increased the persistence of tumor cells at 6 and 24 hours post-administration **(Sup. 4D-E)**. This indicates that both apoptotic fibroblasts and apoptotic tumor cells promote the persistence of live tumor cells over the first 24 hours after arrival in the lung vasculature.

**Figure 2.**
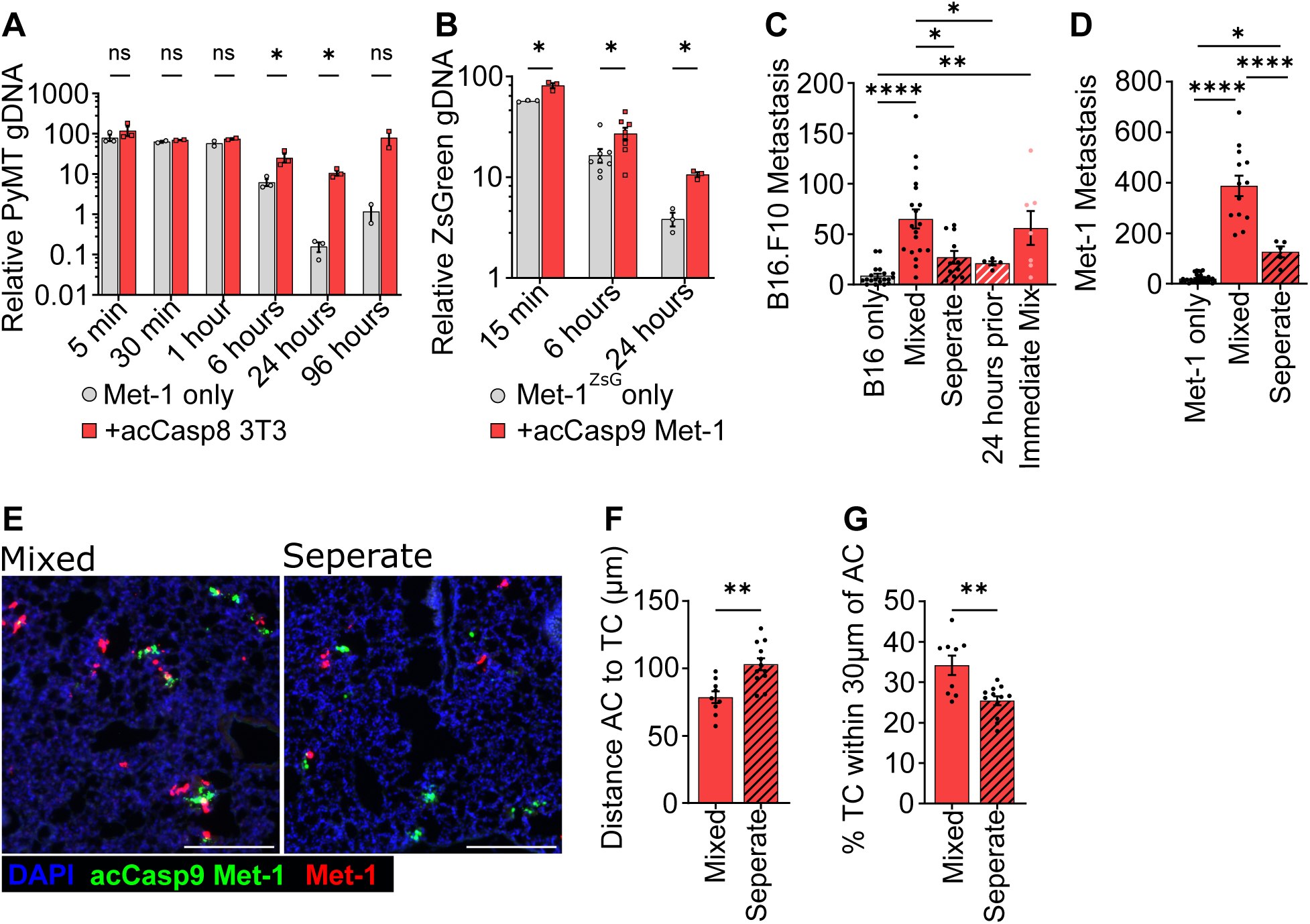
Spatial association of apoptotic cells and tumor cells provides an early survival advantage to tumor cells. Met-1 cells were injected I.V. with acCasp8 3T3 (A) or acCasp9 Met-1 (B) and lung gDNA was harvested at indicated timepoints for qPCR (A-B). Met-1 cells expressed ZsGreen (Met-1^ZsG^) (B). B16.F10 (C) or Met-1 (D-G) cells were injected I.V. alone or in combination with acCasp8 3T3. Tumor cells and apoptotic cells were mixed up to one hour in advance (mixed) or immediately prior and injected in a single bolus, or injected in two sequential injections into opposite tail veins (separate), or apoptotic cells were injected 24 hours prior to tumor cell challenge (C-G). Surface lung metastasis was quantified after 14 days (C-D). Lungs were harvested 1 hour after I.V. for visualization by fluorescent microscopy, acCasp9 Met-1 cells were stained with CFSE (green) and Met-1 cells expressed mCherry (red), scale bar=200µm (E-G). Error bars represent SEM, Dots are biological replicates (A-D) or represent distinct slices of lung tissue taken from n=3-4 biological replicates (E-G), statistical testing is unpaired t-test (A,B,F,G) or Ordinary one-way ANOVA with Tukey’s multiple comparisons test (C-D) AC=Apoptotic Cell, TC=Tumor Cell

The exponential decline in tumor cell persistence over the first 24 hours after I.V. injection is likely a result of those cells undergoing apoptosis themselves. At one hour post-injection, we observed that 40-60% of tumor cells had externalized phosphatidylserine (PS), an indication of early apoptosis **(Sup. 5A-B)**. We observed a reduction in the fraction of PS+ tumor cells at 1-hour post-injection when they were coinjected with apoptotic cells **(Sup. 5C)**. Together, these data suggest that apoptotic cells improve CTC survival by protecting tumor cells during this critical window of lung seeding.

We sought to address whether temporal or spatial separation of apoptotic cells from tumor cells affected metastatic burden by injecting apoptotic cells 24 hours prior to tumor cell challenge, or sequentially into the opposite tail vein from tumor cells. Pre-injecting apoptotic cells 24 hours prior had no effect on metastatic burden in the lung, suggesting that apoptotic cells do not have long-lasting effects that impact CTC survival **(Fig. 2C)**. Injecting apoptotic cells and tumor cells within one minute of each other, but into separate vascular sites, also abrogated the prometastatic effect of apoptotic cells **(Fig. 2C-D)**. These data indicate that apoptotic cells must enter circulation alongside viable tumor cells in order to have a prometastatic effect.

Tumor cells physically associated in clusters with other cells have been shown to have a metastatic advantage over single tumor cells (14,15,40). Given that the murine lung contains over 1km of total vasculature (41), we examined whether injecting apoptotic cells and tumor cells in separate veins reduced their physical association within the lung. When injected together, apoptotic cells were often localized in close proximity to tumor cells within the lung **(Fig. 2E)**. When injected separately into opposite tail veins we observed a reduced percentage of tumor cells in very close proximity to apoptotic cells and an increased average distance between them **(Fig. 2E-G)**. Injecting tumor cells and apoptotic cells in separate tail veins rather than mixed together did not impact the number of tumor or apoptotic cells that seeded the lung. **(Sup. 6A-B)**.

Given these findings, we considered the possibility that tumor cells might receive pro-survival signals from apoptotic cells during the co-incubation period prior to injection. However, we found that the duration of *ex vivo* contact between apoptotic cells and tumor cells did not influence their capacity to enhance metastasis. Combining apoptotic and tumor cells immediately prior to injection or up to 1 hour in advance resulted in comparable enhancement of metastasis by apoptotic cells, suggesting that mixing tumor cells and apoptotic cells does not result in pro-metastatic *ex vivo* cell-cell signaling interactions **(Fig. 2C)**.

Together, these findings indicate that apoptotic cells promote tumor cell survival following initial seeding of the lung vasculature, and suggest that close spatial association between tumor cells and apoptotic cells *in vivo* is required for apoptotic cells to exert these effects. The spatial and temporal dynamics of apoptotic cell enhancement of metastasis led us to hypothesize that apoptotic cells were helping to establish an early protective microenvironment around CTCs.

### Apoptotic cells promote platelet aggregation on tumor cells

Platelets play an essential role in establishing the early metastatic niche and are critical for the survival of tumor cells in circulation (3–5,10). Considering that apoptotic cells promote metastasis by improving CTC survival and that this effect requires their spatial association with tumor cells, we hypothesized that apoptotic cells within the metastatic niche may promote platelet-tumor cell interactions. To examine this, we imaged platelet clots in the lung following injection of tumor cells alone or tumor cells mixed with apoptotic cells. We observed the rapid formation of platelet clots in the lungs after I.V. injection and noticed the clots were often associated with tumor cells and apoptotic cells (**Fig. 3A)**. In mice that received tumor cells plus apoptotic cells we observed significantly more platelet clots as well as a larger average size of platelet clots at 15 minutes post-injection **(Fig. 3B-C).** Platelet clots formed rapidly after tumor cell injection and resolved by 24 hours **(Fig. 3B-C).** Clots that formed around an apoptotic cell were larger than those around a tumor cell, while clots containing both a tumor cell and an apoptotic cell were the largest (**Fig. 3D).** A higher percentage of tumor cells were found in platelet clots in mice coinjected with apoptotic cells than in mice receiving only tumor cells. This additional fraction of tumor cells were localized with apoptotic cells within these clots (**Fig. 3E**). This indicates that apoptotic cells enhance platelet aggregation on tumor cells.

**Figure 3.**
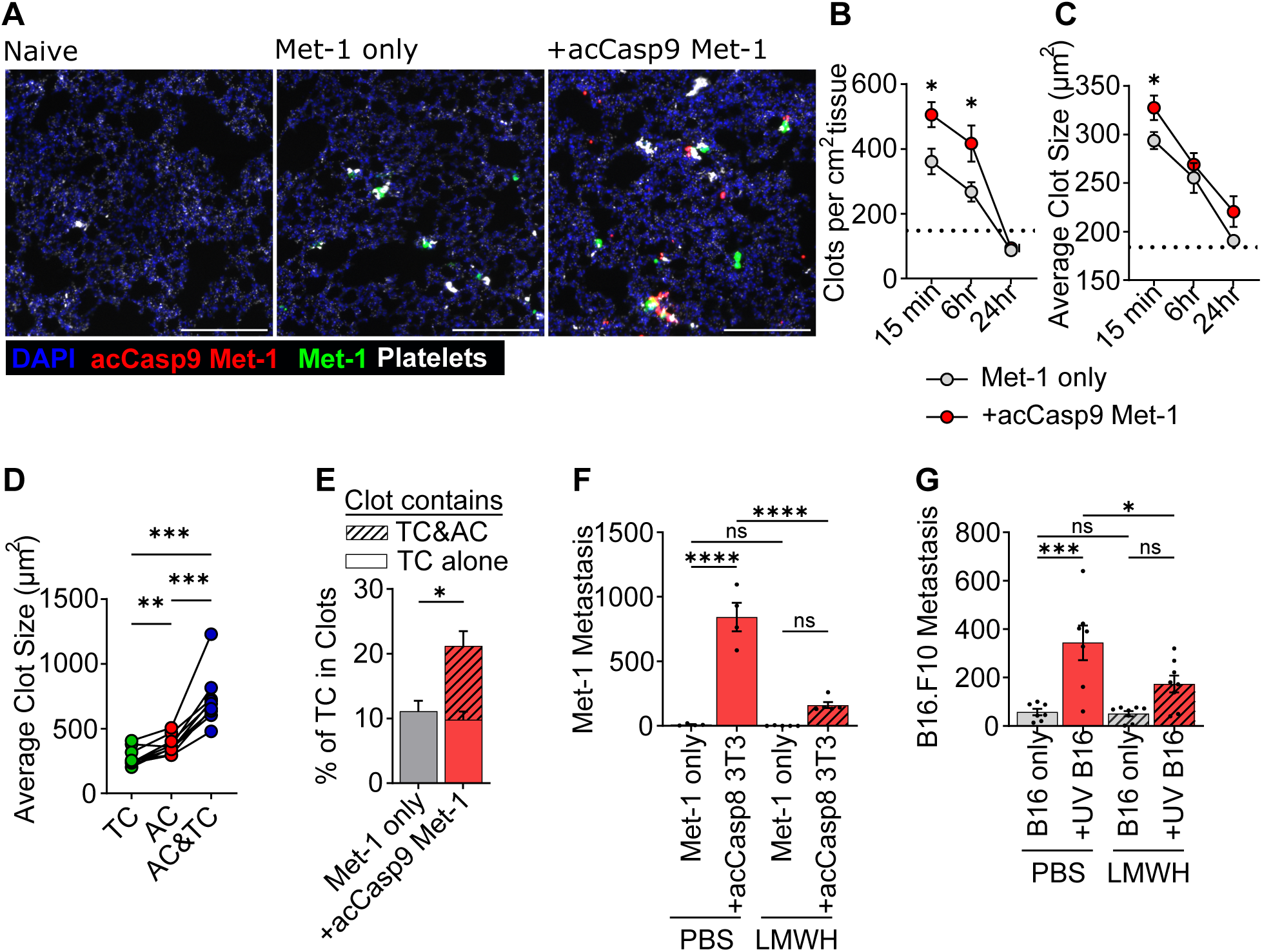
Apoptotic cells promote platelet aggregation on tumor cells and the effect of apoptotic cells on metastasis depends on coagulation. Met-1 cells were injected I.V. with acCasp9 Met-1 (A-E) or acCasp8 3T3 (F). B16.F10 cells were injected I.V. with UV irradiated B16.F10 cells (G). Low-molecular-weight heparin (LMWH) was administered s.c. at −4, 0, and 24 hours in relation to I.V. tumor cell injection. (F-G) Lungs were harvested at 15 min (A,D,E) or indicated timepoints (B,C) and processed for fluorescent imaging. Surface lung metastasis quantified after 14 days (F-G). Met-1 tumor cells expressed ZsGreen (green), apoptotic acCasp9 Met-1 expressed mCherry (red), platelets are stained with αGPIbβ (white), scale bar=200µm (A-E). Dotted line is the mean of naive lung controls (B-C). Average size of clots that contained either an Apoptotic cell (AC), Tumor cell (TC) or an AC and TC was quantified, lines connect values from individual lung sections (D). Striped portion of bar is the % of tumor cells in a clot that also contains an apoptotic cell (E). Error bars represent SEM, dots are multiple slices of lung from n=3-4 biological replicates (B-E) or dots are biological replicates (F-G). Analysis with unpaired T-tests (B,C,E) or by ordinary one-way ANOVA with Tukey’s multiple comparisons test (D,F-G)

We next sought to address whether coagulation was required for the pro-metastatic effects of apoptotic cells. To do this we treated mice with the anti-coagulant Low Molecular Weight Heparin (LMWH) prior to I.V. injection of tumor cells alone or tumor cells and apoptotic cells. In both Met-1 and B16.F10 metastasis models we observed that treating mice with LMWH reduced the effect of apoptotic cells on metastasis **(Fig. 3F-G)**. This suggests that coagulation is essential for apoptotic cells to promote metastasis.

### Apoptotic cells have enhanced procoagulant activity

Given our *in vivo* data indicating a role for coagulation in the promotion of metastasis by apoptotic cells, we sought to directly assess the pro-coagulant activity of apoptotic cells *in vitro*. To do this, we developed an assay based on the prothrombin time coagulation test, a test which measures how fast a clot forms within plasma (42). We quantified how rapidly live or apoptotic cells formed a fibrin clot in citrated, platelet-poor plasma by measuring the change in absorbance after addition of Ca^2**+**^ **(Sup. 7)**. A lower time to clot formation indicates higher pro-coagulant activity. When Met-1 or 3T3 cells were stimulated to undergo apoptosis, the apoptotic cells were faster at inducing a fibrin clot compared to their live cell counterparts, as evidenced by a lower time to reach V_max_ **(Fig. 4A)**. Notably, 3T3 cells were overall faster at initiating coagulation compared with Met-1 cells, mirroring the more robust promotion of metastasis observed upon coinjection of apoptotic 3T3 cells as compared to apoptotic Met-1 cells *in vivo*.

**Figure 4.**
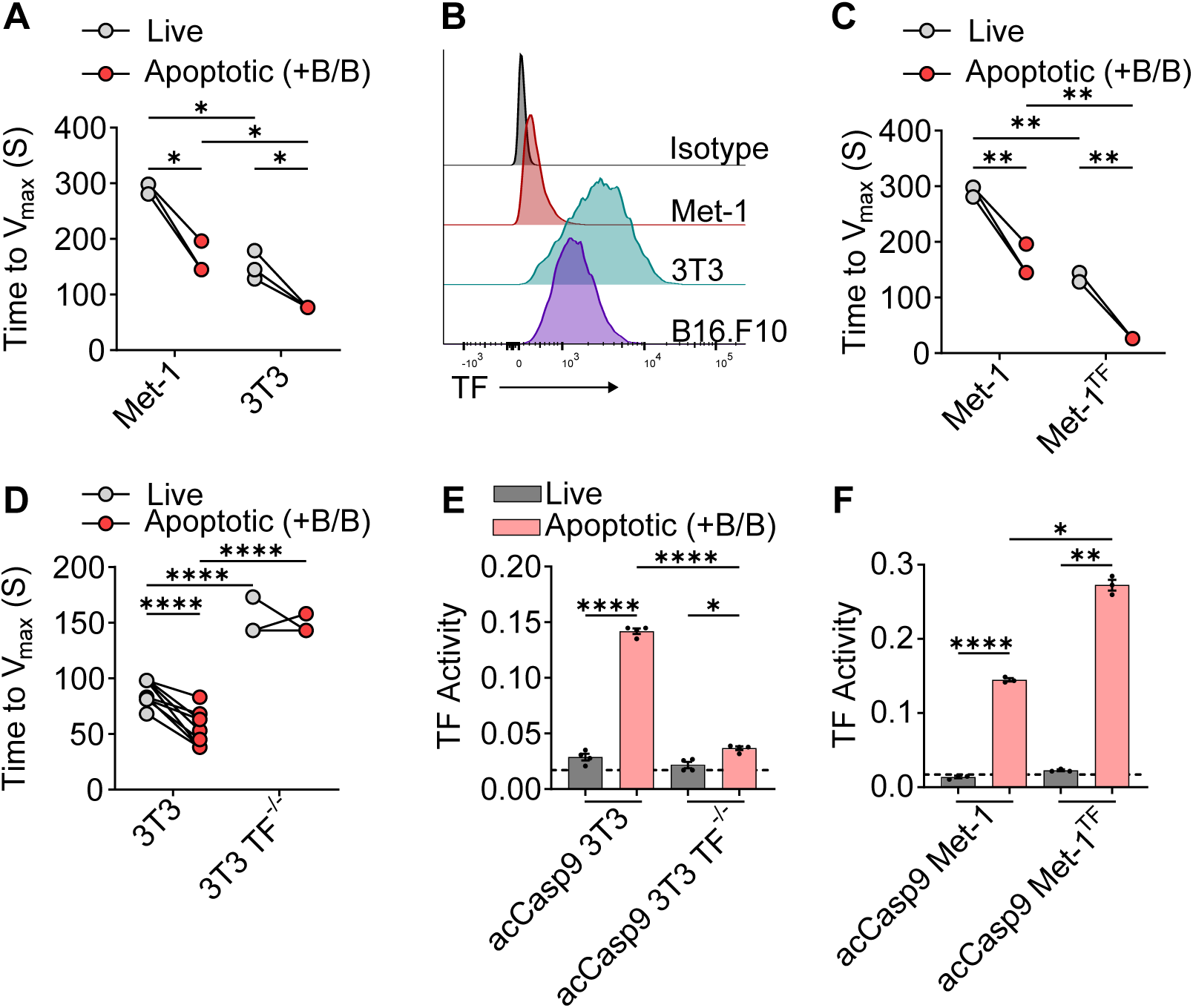
Apoptotic cells have enhanced procoagulant activity. Met-1 or 3T3 expressing acCasp9 were left untreated or treated with B/B for 2 hours (A,C-F). Time to fibrin clot formation was measured as the maximum change in absorbance (Time to V_max_) after adding Ca^2+^ to citrated mouse plasma containing live or apoptotic cells, lower values indicate more rapid clot formation (A,C,D). TF-PE fluorescent intensity was measured by flow cytometry (B). TF activity was measured as the absorbance at 30 minutes after addition of live or apoptotic cells to mouse FVII, human FX, and chromogenic FXa substrate (E,F). Circles connected by lines are matched serum samples and statistical analysis is 2way ANOVA with Šidák multiple comparison test (A,C,D). Dotted line represents the absorbance of a control well containing all assay components without cells added, statistical analysis is Welch’s ANOVA with Dunnett’s multiple comparisons test, error bars represent SEM (E-F).

Plasma contact with the cell surface protein TF initiates the extrinsic coagulation cascade (43). High TF expression on tumor cells has been linked with cancer-associated thrombosis and metastatic progression (11,44–46). We wondered if the difference in coagulation rates of Met-1 vs. 3T3 cells could be due to differential expression of TF. Consistent with this possibility, Met-1 cells exhibited substantially lower expression of TF than 3T3 or B16.F10 cells **(Fig. 4B).** We therefore generated TF knockout 3T3 cells (3T3 TF^-/-^) or TF overexpressing Met-1 cells (Met-1^TF^) to test whether TF was responsible for their procoagulant activity. Indeed, we observed that the rate of fibrin clot formation induced by each cell type correlated with TF expression. Overexpressing TF on Met-1 cells increased their rate of plasma clot formation **(Fig 4C)** while ablating TF on 3T3 cells reduced their rate of plasma clot formation. Notably, ablation of TF on 3T3 cells abrogated the higher procoagulant activity of apoptotic cells compared to live cells **(Fig 4D)**. This indicates that the speed of clot formation by apoptotic cells depends on TF-mediated coagulation cascades.

TF activates coagulation by forming a complex with Factor VIIa, which catalyzes the conversion of Factor X to Factor Xa. However, TF often exists in an inactive state, termed encrypted TF, on resting cells, which prevents efficient catalysis of this reaction (47). While the exact mechanisms controlling TF decryption remain incompletely defined, various cell perturbations, including those that induce PS externalization, can reveal full TF procoagulant activity (47). We thus directly examined TF activity on live vs. apoptotic cells by assaying the rate of Factor Xa generation in the presence of Factor VII and Factor X. TF on live cells was largely unable to generate Factor Xa irrespective of its expression level suggesting that TF is in the encrypted state on live Met-1 and 3T3 cells. Inducing apoptosis significantly increased TF activity on these cells **(Fig 4E-F).** The magnitude of TF activity after induction of apoptosis was dependent on the level of TF expression. Overexpressing TF on acCasp9 Met-1 or knocking it out on acCasp9 3T3 led to a corresponding increase or decrease of TF activity respectively **(Fig 4E-F).** Together, these *in vitro* data suggest that inducing apoptosis leads to TF decryption, resulting in enhanced procoagulant of apoptotic cells.

### Phosphatidylserine exposure promotes the coagulant activity of apoptotic cells

PS externalization is an important mediator of thrombosis that occurs during platelet activation and endothelial damage and contributes to other pathological thrombotic events. PS externalization has been described as a mechanism for decrypting TF pro-coagulant activity as well as providing a negatively charged surface for assembly of the FXa-Va-prothrombinase coagulation enzyme complex (47–50). PS externalization is also a canonical feature of the apoptotic plasma membrane; we therefore investigated whether the procoagulant activity of apoptotic cells was PS-dependent. We employed two distinct strategies to target PS. First, to physically block PS exposed on the outer membrane, we treated cells with saturating concentrations of the PS binding protein Annexin V (AxV). We also generated apoptotic cells that do not externalize PS upon caspase activation by deleting the caspase-activated scramblase XKR8 and expressing a caspase-resistant version of the flippase ATP11c (51) (acCasp9 3T3 Flip^mut^) **(Sup. 8).** Blocking PS with AxV completely abrogated the ability of TF to generate Factor Xa in the presence of Factor VII and Factor X **(Fig. 5A).** acCasp9 3T3 Flip^mut^ cells unable to externalize PS also lacked TF activity after induction of apoptosis. **(Fig. 5A)** These data suggest that TF decryption is dependent on PS externalization during apoptosis.

**Figure 5.**
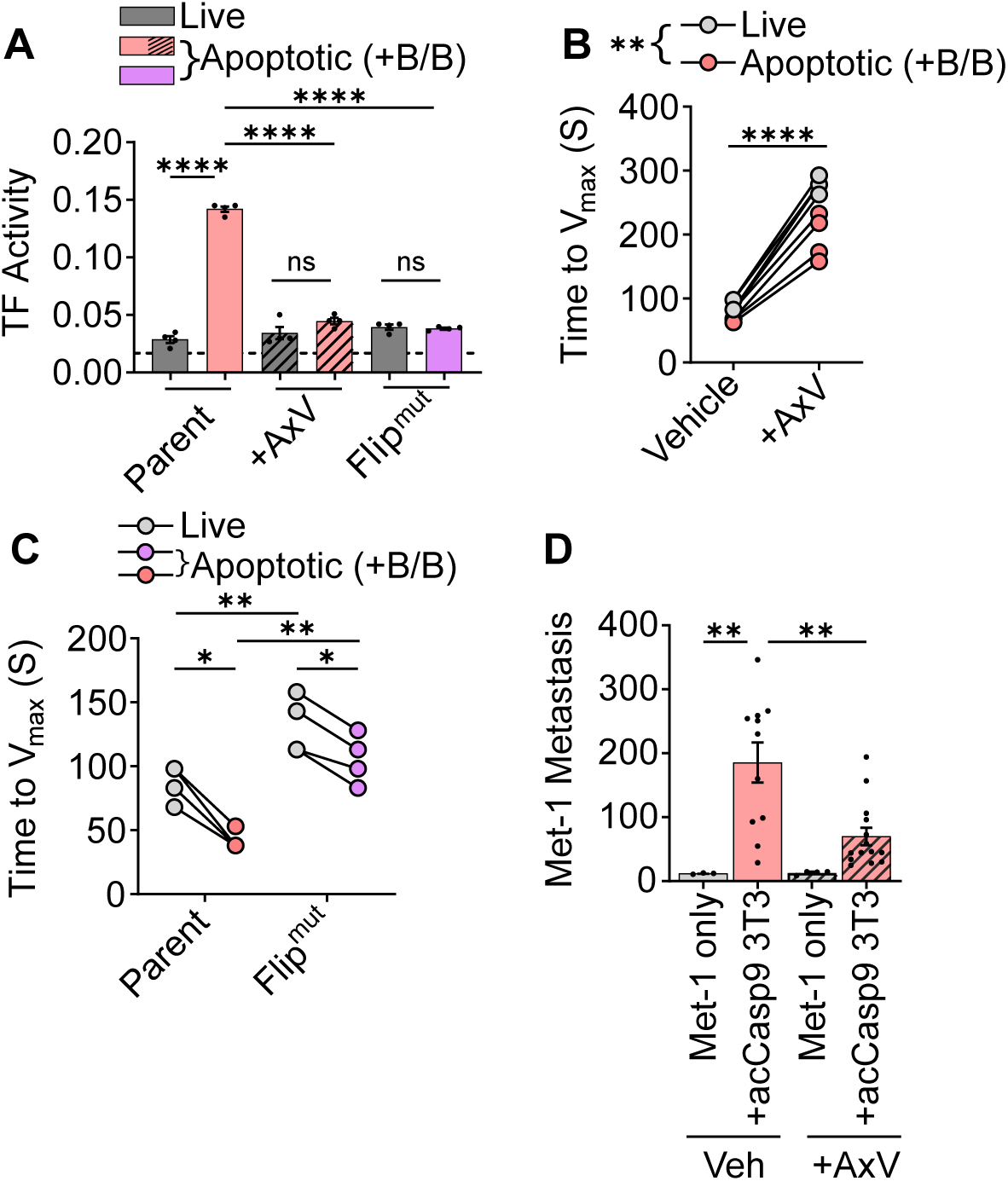
Phosphatidylserine exposure promotes the coagulant activity of apoptotic cells. acCasp9 3T3 were left untreated or treated with B/B for 2 hours (A-C) or 15 minutes (D). TF activity was measured as the absorbance at 30 minutes after addition of live or apoptotic cells to mouse FVII, human FX, and chromogenic FXa substrate (A). Time to fibrin clot formation was measured as the maximum change in absorbance (Time to V_max_) after adding Ca^2+^ to citrated mouse plasma containing live or apoptotic cells, lower values indicate more rapid clot formation (B,C). Annexin V (AxV) was added to cells in Ca^2+^ containing buffer and incubated on ice for 30 minutes (A,B,D). Surface lung metastasis was quantified at 14 days post I.V. injection (D). Dotted line represents the absorbance of a control well containing all assay components without cells added (A). Statistical analysis is Welch’s ANOVA with Dunnett’s multiple comparisons test, error bars represent SEM (A,D). Circles connected by lines are matched serum samples and statistical analysis is 2way ANOVA with Šidák multiple comparison test (B,C).

To examine whether the full procoagulant activity of apoptotic cells was dependent on PS externalization, we next tested these PS blocking strategies in the plasma clot assay. AxV treatment significantly reduced the plasma coagulation rate, however, apoptotic cells treated with AxV still induced clotting faster than live cells treated with AxV **(Fig. 5B).** acCasp9 3T3 Flip^mut^ cells also retained an increased plasma coagulation rate upon caspase activation, despite having reduced overall procoagulant activity compared to parental cells **(Fig 5C)**. Together this indicates that while TF decryption and maximal procoagulant activity on apoptotic cells is dependent on PS, apoptotic cells have residual, PS-independent procoagulant activity.

We next tested whether blocking PS resulted in reduced metastasis *in vivo*. We found that treatment of apoptotic cells with AxV significantly reduced their ability to promote metastasis **(Fig. 5D)**. The early benefit of apoptotic cells on tumor cell survival at 6 and 24 hours was also reduced by treating cells with AxV **(Sup. 9A-B)**. This suggests that PS externalization on apoptotic cells benefits early CTC survival and metastasis.

### TF expression is required for apoptotic cells to promote clotting *in vivo* and enhance metastasis

Given the differential expression levels of TF on Met-1 vs. 3T3 cells **(Fig. 4B)** and the corresponding difference in magnitude of their effect on coagulation and metastasis **(Fig. 1 and Fig. 4a)**, we next sought to directly assess the role of TF in the promotion of metastasis by apoptotic cells. TF overexpression on apoptotic Met-1 cells increased their effect on metastasis, to a level comparable to apoptotic 3T3 **(Fig 6A-B)**. Conversely, TF knockout reduced the effect of apoptotic 3T3s to a level comparable to that observed with apoptotic Met-1 **(Fig 6C-D)**. The relative tumor cell survival at 24 hours was also dependent on TF expression by apoptotic cells **(Sup. 9C-D)**. This suggests that TF expression and the procoagulant activity of apoptotic cells determines the magnitude of effect apoptotic cells have on promoting metastasis.

**Figure 6.**
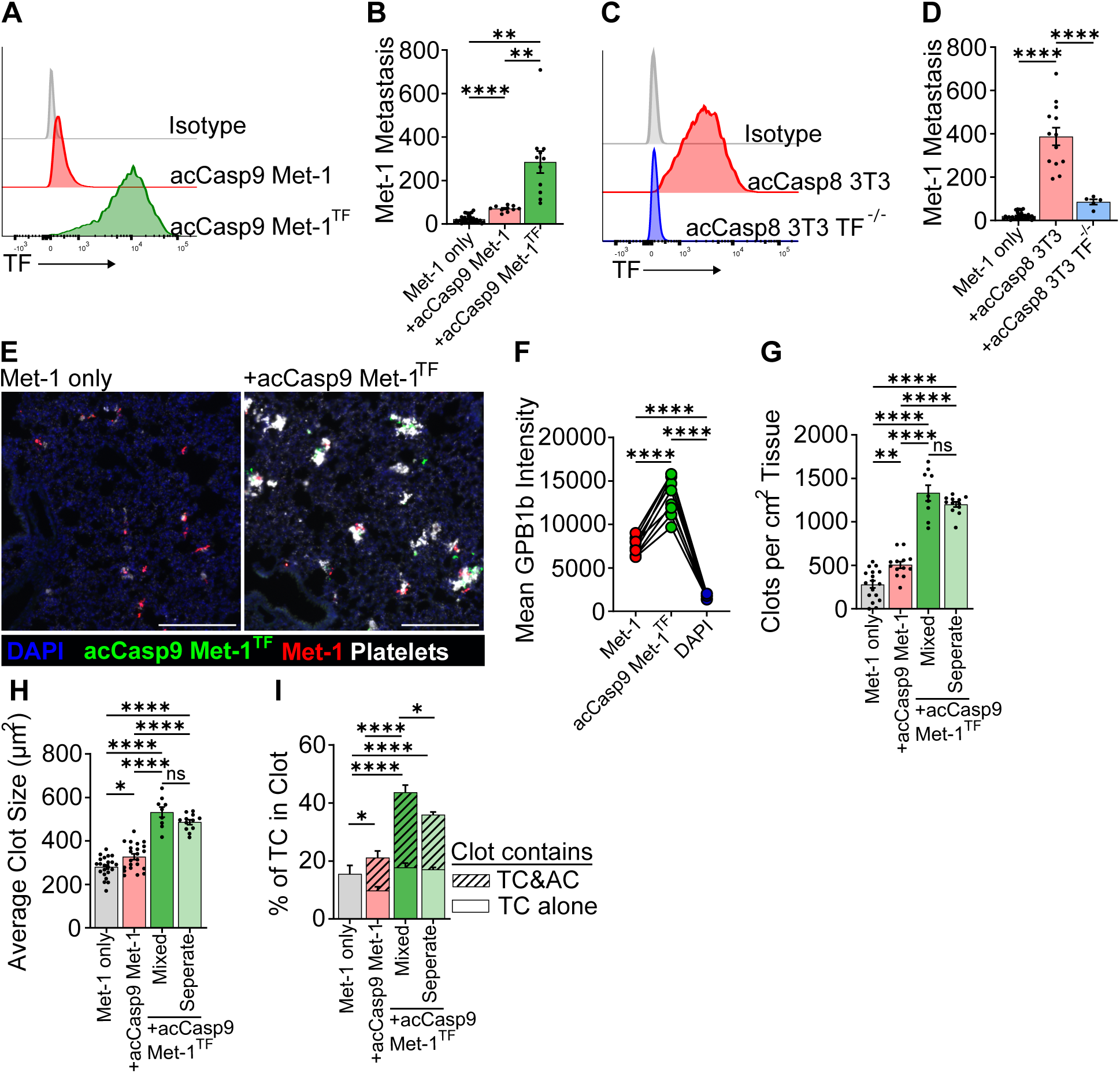
Tissue Factor is responsible for the magnitude of apoptotic cell enhanced metastasis and platelet clotting in the lung. TF was overexpressed on acCasp9 Met-1 cells or knocked out on acCasp8 3T3 cells. TF-PE fluorescent intensity was measured by flow cytometry on live cells (A,C). acCasp9 Met-1 (B,E-I) or acCasp8 3T3 (D) were treated with B/B for 15 minutes. Surface lung metastasis was quantified at 14 days post I.V. injection (B,D). Lungs were harvested for fluorescent microscopy 1 hour after I.V. injection, Met-1 and acCasp9 Met-1^TF^ cells were mixed up to one hour in advance and injected in a single bolus (mixed), or injected in two sequential injections into opposite tail veins (separate), acCasp9 Met-1^TF^ cells were stained with CFSE (green), Met-1 cells expressed mCherry (red), platelets are stained with αGP1bβ (white), scale bar=200µm (E-I). Mean GPB1b intensity on Met-1 tumor cells, acCasp9 Met-1^TF^ apoptotic cells, or across all DAPI+ tissue was quantified from mice that received Met-1 mixed with acCasp9 Met-1^TF^, lines connect values from individual lung sections (F). Striped portion of bar is the % of tumor cells in a clot that also contains an apoptotic cell (I). Dots are biological replicates (B,D) or multiple slices of lung from n=3-4 biological replicates (F-I). Statistical testing is Ordinary one-way ANOVA with Tu key’s multiple comparisons test. Error bars represent SEM. AC=Apoptotic Cell, TC=Tumor Cell

Given our previous observation that apoptotic cells promoted platelet aggregation at metastatic sites **(Fig 3)**, we next assessed the role of TF on apoptotic cells in this process. We found that the lungs of mice coinjected with TF expressing apoptotic cells contained large platelet clots **(Fig 6E)**. The intensity of platelet staining associated with apoptotic Met-1 cells overexpressing TF was significantly higher than that observed on unmanipulated Met-1 tumor cells **(Fig 6F)**. Increasing TF expression on apoptotic cells significantly increased the number and size of clots found in the lung **(Fig 6G-H)**. This suggests that expression of TF on apoptotic cells drives robust platelet activation during vascular dissemination.

Finally, we assessed the role of TF-overexpressing apoptotic cells in promoting aggregation of platelets directly on Met-1 tumor cells with low TF expression. To do this, we quantified the percent of tumor cells within platelet clots and enumerated clots that contained only a tumor cell and those that contained both a tumor cell and an apoptotic cell. Compared to the small increase in tumor cell localization to clots when coinjected with TF-low apoptotic cells, TF-high apoptotic cells led to a substantial increase in the percent of viable Met-1 tumor cells found within clots. This increase was reflected in the portion of viable tumor cells localized to a clot that also contained a TF-expressing apoptotic cell, suggesting that TF expression on apoptotic cells can act *in trans* to promote platelet aggregation on tumor cells that have low TF expression. **(Fig. 6I)**

We also examined whether injecting apoptotic cells separately from tumor cells, into opposite tail veins, had an effect on the localization of tumor cells and apoptotic cells within clots. While the overall size and number of clots was not affected by injecting apoptotic cells and tumor cells separately, the frequency of tumor cells localized to a clot with an apoptotic cell was decreased **(Fig. 6G-I).** This indicates that when apoptotic cells and tumor cells are injected separately, they are less likely to localize together within clots. Because injecting tumor cells and apoptotic cells in separate veins reduces their effect on metastasis **(Fig. 2C-D)** and the effects of apoptotic cells are dependent on their ability to trigger coagulation through TF **(Fig. 4,6)**, we postulate that tumor cells localized together in a clot with an apoptotic cell are more likely to survive and form a metastatic nodule. Together, these findings identify a previously unappreciated role for apoptotic cells in promoting metastasis, and suggest that TF expression by apoptotic tumor cells is a key contributor to this process.

## DISCUSSION

Killing tumor cells by apoptosis is a central goal of most tumor therapies. Counterintuitively however, an expanding body of evidence suggests that the presence of apoptotic cells, from either tumor or stromal origin, may correlate with increased tumor growth and negative clinical outcome. While apoptotic cell death within tumors is known to promote tumor cell proliferation, angiogenesis, and pro-tumor macrophage polarization (52–54), a putative role of apoptotic cells throughout the metastatic cascade has not been well defined. Here we report a striking role for apoptotic cells in facilitating metastasis by supporting early CTC survival. We find that the effect of apoptotic cells on metastasis depends on the pro-coagulant activity of TF, which is induced upon PS exposure on apoptotic membranes.

In imaging the early seeding of CTCs in the lungs, we observed that apoptotic cells promote platelet aggregation, and that the co-localization of apoptotic cells with tumor cells thereby supported CTC survival. Platelet aggregation on tumor cells is important for shielding tumor cells from NK cell cytotoxicity and reducing shear stress, two significant contributors to early CTC demise (3–8); our findings suggest that the presence of apoptotic cells offers a mechanism for CTCs to activate coagulation to survive these pressures. Extrinsic coagulation mediated by TF is rapid, and intravascular aggregation of CTCs is rare (14); thus, when apoptotic and tumor cells do not enter circulation together—modeled herein as separate injections into opposing tail veins—apoptotic cell-induced coagulation would not occur in proximity to a tumor cell, thus abrogating the metastatic benefit conferred by apoptotic cells. CTC clusters have been shown to metastasize significantly more effectively than individual CTCs. CTC clusters disseminate collectively, entering circulation as a single unit (14,15), and thus represent a physiological context in which an apoptotic cell could be localized with a tumor cell in circulation. While, various pro-metastatic characteristics of CTC clusters have been defined such as stemness, apoptosis resistance, and immune escape (13,16), the presence of apoptotic cells within clusters has not been investigated. An intriguing possibility for one metastatic advantage conferred by CTC clusters is that a portion of cells in clusters could undergo apoptosis, thus supporting the survival of other cells within the cluster. Technological advances in detecting and analyzing CTC clusters could shed light on the clinical and physiological relevance of apoptotic cells as a component of CTC clusters.

Mechanistically, we find that apoptotic cells initiate robust coagulation through PS mediated TF decryption. An important role of TF expression on CTCs that successfully metastasize has been extensively described (45,55); however, our work suggests that TF mediated coagulation can also act *in trans* to support CTC survival. Thus, tumor cells with low TF expression, such as Met-1 cells, may depend on apoptosis of other cells as a source of TF activity to facilitate coagulation during vascular dissemination. Fibroblasts have been observed in association with CTCs (56–58) and are a potential source of TF mediated coagulation during metastasis that may be particularly relevant for the survival of TF-low tumor cells in circulation.

Our findings bolster the concept that apoptotic cells are an important procoagulant mechanism in various health and disease settings (59). TF activity regulated through phosphatidylserine exposure has been described in other disease contexts such as endotoxemia (60–62) and vessel injury(63,64). Most relevant to our model is the consideration of apoptotic cells as a contributing factor to cancer-associated thrombosis (65–68). Interestingly, the risk of venous thromboembolism increases following chemotherapy (69–71), a regimen that causes massive apoptosis throughout the body. While chemotherapy has numerous effects beyond induction of apoptosis, our data clearly demonstrate that direct activation of caspases in both tumor and non-tumor cells is a source of procoagulant activity, suggesting that apoptotic cells may be an important contributor to hypercoagulation following chemotherapy.

While our work establishes apoptotic cells as a source of TF and PS-induced coagulation that supports CTC survival in the vasculature, other sources of TF and PS exist and the differential effects of these unique activators of coagulation throughout the metastatic cascade remains an open question. Extracellular vesicles are a source of procoagulant activity and can exhibit pro-metastatic properties (68,72,73). Apoptosis results in membrane blebbing and release of extracellular apoptotic vesicles, but whether there are distinct outcomes depending on whether EVs are derived from live or apoptotic cells is not clear. Tumor cells themselves have elevated surface PS and targeting PS as a therapeutic strategy has been promising (30,74–79). We observed PS externalization on tumor cells in circulation, and interpreted this observation as an early marker of apoptosis. However, it is also possible that deformation and shear stress in circulation results in Ca^2+^ flux which could activate the TMEM family of PS scramblases (80), resulting in PS externalization on live cells. Technical limitations in visualizing externalized PS *in vivo* prevent observation of PS externalization in combination with platelet aggregation, so we cannot yet delineate the potential contributions of PS exposure on live cells to coagulation *in vivo*. Necrotic cells also have procoagulant activity (61,81–83), yet these cells do not promote metastasis. Although the procoagulant activity of apoptotic cells is required, additional features of apoptotic cells could be responsible for the differential effect of apoptotic and necrotic cells on metastasis. While numerous possible sources of procoagulant activity exist during metastasis, elucidating the differential subsequent effects of apoptotic cells compared to other sources of coagulation throughout the metastatic cascade is warranted.

While our data highlight the key role of apoptotic cells in initiating coagulation to support metastasis, we are intrigued by the possibility that additional mechanisms exist by which apoptotic cells influence the metastatic process. Apoptotic cells alter their surrounding microenvironment in profound ways. Molecules such as fractalkine, lactotransferrin, and prostaglandin E_2_ are released by apoptotic cells to recruit macrophages and can also act on tumor cells to promote survival and proliferation (53,84–87). Upon caspase activation, apoptotic cells release various anti-inflammatory metabolites from pannexin-1 channels (88). Interestingly, ATP released through pannexin-1 was demonstrated to have pro-survival signaling in an autocrine manner on CTCs (89). In the context of a CTC cluster, these various signaling molecules released from apoptotic cells could act in a paracrine fashion to support nearby tumor cells. Furthermore, the interactions between apoptotic cells and macrophages has been extensively characterized as leading to tissue-repair macrophage phenotypes (29). Previous work has demonstrated that efferocytosis of apoptotic cells within tumors can polarize macrophages, which produce TGF-β and other wound-healing cytokines to facilitate metastatic dissemination (27). CTCs release microparticles upon early arrival in the lung vasculature that are sampled by myeloid cells and guide their differentiation (90). We observe rapid uptake of apoptotic cells by phagocytes in the lungs; however, whether the interaction between apoptotic cells and phagocytes contributes to the differentiation of metastasis associated macrophages is an open question. The downstream effects of apoptotic cells beyond their most proximal role in inducing coagulation during hematogenous dissemination is an interesting area for future investigation.

Decades of work has been aimed at finding mechanisms to kill tumor cells, primarily through inducing apoptosis. Our work supports a growing body of evidence that apoptosis in the context of cancer can have adverse pro-tumor effects. Eliminating tumor cells, primarily through apoptotic cell death, will remain the primary goal of cancer therapeutics. However, understanding the subsequent effects of apoptosis on cancer progression could inform additional interventions countering the pro-tumor secondary effects of apoptosis. For example, blocking apoptosis by inhibiting caspase activation results in the conversion of apoptosis to an inflammatory form of cell death capable of enhancing anti-tumor immunity (91). Whether this strategy could also reduce metastasis should be addressed. Because we define apoptotic cells as an important source of coagulation and show that LMWH treatment reduces the effect of apoptotic cells in promoting metastasis, targeted anti-coagulants are another possible therapeutic strategy. Temporal administration of anti-coagulants or the use of PS or TF blocking agents during times where high levels of apoptosis are expected, such as during cytotoxic therapy, could specifically target coagulation initiated by apoptotic cells. While the use of anti-coagulants has been proposed as a potential therapeutic to reduce metastatic spread (92,93), they are not yet broadly used as anti-metastatic agents. The safety and efficacy of anti-coagulants in patients at risk for cancer-associated thrombosis is being evaluated (94,95); our data suggest than an interesting additional parameter to monitor in these trials could be progression to metastatic disease.

Building upon clinical and experimental evidence that apoptosis can have unintended pro-tumor consequences, this work demonstrates a direct mechanism by which the presence of apoptotic cells during vascular CTC dissemination promotes metastatic disease. By activating coagulation through PS-dependent TF decryption, apoptotic cells support the survival of tumor cells in circulation. These findings emphasize that it is important to consider not only the cells that directly form metastatic tumors, but also the effects of cells that die throughout the metastatic cascade, opening up new avenues for investigation into the process of metastasis and possibilities for therapeutic intervention.

## METHODS

### Sex as a biological variable

Our study examined only female animals because breast cancer metastasis is a disease burden primarily in females. It is unknown whether the findings are relevant for male mice.

### Cell Culture

Cell lines were cultured at 37C with 5% CO2 in Dulbecco’s modification of Eagle medium (DMEM) supplemented with 10% (v/v) fetal bovine serum (FBS), 2 mM l-glutamine, 10 mM Hepes, and 1 mM sodium pyruvate (complete DMEM). Cell lines were passaged every 1-3 days and used for experiments within 2 weeks after thawing from liquid nitrogen stocks. Cells were regularly tested and remained negative for mycoplasma contamination. Adherent cells were detached from plate using Trypsin-EDTA (0.25%). Mouse Embryonic Fibroblasts (MEFs) were derived from day 15.5 embryos of B6/J or FVB/N pregnant mice as previously described(96). MEFs were immortalized by transforming with lentivirus containing SV40 Large T antigen (iMEF). B16.F10 (CRL-6475) and NIH/3T3 (CRL-1658) cells were purchased from ATCC. Met-1 cells were derived from mammary carcinomas in FVB/N-Tg(MMTV-PyVmT) and kindly provided by Dr. Alexander Borowsky (31). Met-1^TF^ cells were generated by lentiviral transformation with pCMV3-C-GFPSpark containing the mouse Tissue Factor ORF (SinoBiological). NIH/3T3 TF^-/-^ cells were generated by lentiviral transformation with pRRL.U6.gRNA.MND.Cas9.2A.Hygro containing the guide RNA sequence CTCGTCTGTGAGGTCGCACT. Flip^mut^ cells were generated by lentiviral transformation with pRRL.U6.gRNA.MND.Cas9.2A.Hygro containing the guide RNA sequence targeting *XKR8*, CGTACTGGACAACGGCCCAC, and pMX.puro containing a mutant of ATP11c with D to A mutations in the three caspase cleavage sites, kindly provided by Dr. Shigekazu Nagata (51).

### Mice

Female C57BL6/J (B6/J) mice and FVB/NJ mice (FVB) were purchased from the Jackson Laboratory and allowed to acclimate at least 1 week before experiment initiation. Experiments were performed on 6-10 week old mice. A *Rag2^-/-^* breeding pair was purchased from the Jackson Laboratory and bred in house for experiments. Mice were housed under specific pathogen–free conditions at the University of Washington. All animals were maintained according to protocols approved by the University of Washington Institutional Animal Care and Use Committee (IACUC), under protocol number 4298-01 (PI: AO).

### Cell Death Induction

The activatable cell death systems have been described extensively previously (32–35). Briefly, NIH/3T3 cell lines were transduced with lentiviral plasmid (pRRL) encoding activatable (“ac”) Caspase-8, Caspase-9 or RIPK3. pSLIK lentiviral vector was used for thyroid response element controlled expression of acCasp9 in B16.F10 and Met-1 cell lines and GSDME in NIH/3T3 cells. pSLIK gene expression was induced by culturing cells in doxycycline (1 µg/ml; Sigma-Aldrich) for 18 hours before harvesting for cell death induction. Programed cell death was induced by incubating cells in complete DMEM with 1 mM B/B homodimerizer (Clontech) for 15 min at 37°C. Cells were then washed with cold phosphate-buffered saline (PBS) before being resuspended in PBS at designated concentrations. Cells killed by freeze/thaw lysis were cycled between liquid nitrogen and a 37°C water bath 3 times. Cells killed by UV irradiation were exposed to 100mj/cm^2^ UV light using a UVP CL-1000 Ultraviolet Crosslinker. % Cell death was monitored using Sytox Green dye (Invitrogen) and Incucyte live cell imaging (Sartorius). Phosphatidylserine exposure was analyzed on a flow cytometer after staining cells with Fluorescent conjugated Annexin V (Invitrogen) and Propidium Iodide (Ebioscience) in Annexin Binding Buffer.

### Intravenous Metastasis

1.5×10^5^ Met-1 tumor cells or 7×10^4^ B16.F10 cells were suspended in PBS with an equivalent number of apoptotic cells. 100µL of this mixture was injected intravenously into the tail vein of mice after warming under a heat lamp for 5 minutes. B16.F10 metastasis were quantified 14 days after by counting black tumor nodules on the surface of all lung lobes, Met-1 metastasis was quantified by staining lung tissue with 15% India Ink intratrachealy, then incubating lungs in Fekete’s solution for 24 hours at 4°C before counting white tumor nodules on the surface of all lung lobes. For separate tail vein experiments apoptotic cells and tumor cells were suspended in PBS and 100µL of each cell type was injected into the left and right tail veins respectively. For phosphatidylserine blocking, 25µg/mL purified Annexin V (Biolegend) was added to cells in HBSS+Ca^2+^+Mg^2+^.

### Quantification of injected cell genomic DNA in Lung

The left lobe was dissected, rinsed in PBS, and placed in 1mL DNAzol (Invitrogen). The tissue was homogenized using Precellys Tissue Homogenizer with ceramic beads. Genomic DNA was isolated from homogenate according to the DNAzol reagent manual. The concentration of genomic DNA isolated was quantified using spectrophotometer and concentrations were normalized by diluting samples in nuclease-free water. Quantitative Taqman probe based PCR was used to quantify the PyMT transgene or other ectopically expressed transgenes (ZsGreen, GFP, mCherry). Primer/Probe sequences can be found in supplemental table 1. The relative abundance of target genes was calculated as 2^-ΔΔCT^ where ΔCT=CT^Target^-CT^ptger2^ and ΔΔCT= ΔCT^sample^-Average ΔCT^initial^.

### Heparin Treatment

200µg of low molecular weight heparin, Enoxaparin sodium (Sigma Aldrich) dissolved in 200µL PBS was injected subcutaneously into the flank of mice at hours −4, 0 and 24 relative to I.V. tumor cell injection.

### NK cell depletion

Mice were administered 200µg anti-NK1.1 clone PK136 (Bio X Cell) or isotype mouse IgG2a clone C1.18.4 (Bio X Cell) in 200 µL PBS intraperitoneally at 72 and 24 hours prior to I.V. metastasis challenge.

### Flow Cytometry

Lungs were rinsed in PBS, roughly dissociated with scissors, then incubated with 50µg/mL Liberase TM (Sigma), 250µg/mL DNAse I (Sigma) in HBSS with Ca^2+^&Mg^2+^ for 35 minutes at 37C with gentle agitation. Tissue was homogenized with GentleMACS lung dissociation (Miltenyi Biotec) and strained through 70µm cell strainers. Single cell suspensions were stained with fluorochrome conjugated antibodies in PBS with 3% FBS +0.05% NaAzide. Antibodies used are CD45 clone 30-F11 (Biolegend), Tissue Factor polyclonal (R&D Systems), CD3 clone 145-2C11 (BD Bioscience), NKp46 clone PK136 (Biolegend). Fluorescent conjugated Annexin V (Invitrogen) and Propidium Iodide (Ebioscience) staining was added to cells suspended in Annexin Binding Buffer prior to running on either a CantoRUO or Symphony flow cytometer (BD Biosciences). Data was analyzed with FlowJo software (TreeStar).

### Immunofluorescent imaging

Mouse lung tissue was fixed in a 1:3 dilution of Cytofix fixation buffer (BD) in PBS for 18 hours, washed with PBS twice, then embedded in OCT medium and frozen. In some experiments, cells were stained with CFSE (Invitrogen) prior to injection. 20µm slices were prepared with a Cyrostat on glass microscope slides and stained with primary and secondary antibodies in 1%BSA in PBT (PBS with 0.1% Triton X-1000) after blocking with 10% BSA in PBT. Antibodies used were anti-GPIbβ conjugated to Dylight 649 (X649, Emfret analytics); and rabbit anti-mCherry polyclonal (Rockland). Slides were mounted with Aqua-mount (Epredia) and images were acquired on a Leica DMI6000 inverted microscope with 10X LWD dry objective and Leica DFC365 FX CCD camera. LASX software was used for image acquisition and mosaic tiling.

### Image analysis

ImageJ was used to quantify platelet clots, tumor cell numbers, and apoptotic cell numbers. Positive staining was defined using manual thresholding. Clots, tumor cells, and apoptotic cells were defined as stain positive objects >200µm^2^. QuPath was used to quantify spatial parameters such as colocalization of cell within clots and distances between cells using manual thresholding to define objects.

### Plasma Coagulation Assay

Blood was drawn from the left ventricle into syringes containing citrate-dextrose solution (Sigma-Aldrich) at a dilution of 9:1. Platelet-poor plasma was isolated after two sequential centrifugations at 5,000xg for 10 minutes. Plasma was stored at −80 and thawed only once before use in assay. All reagents were brought to 37°C for assay. 50µL citrated platelet-poor plasma was aliquoted into wells of a 96 well polystyrene plate. 100,000 cells were added to wells in 50µL PBS. 50µL of 25mM CaCl_2_ was added to allow the coagulation reaction to occur. Absorbance at 405nm was measured every 15 seconds by the Syngery HT microplate reader at 37°C with a 2 second shaking step in between each read. The time to the maximum change in absorbance was calculated (Time to V_max_).

### Tissue Factor activity assay

Live cells or cells stimulated to undergo apoptosis (20,000 per well) were added to a 96 well plate. Hepes buffered saline with BSA (1mg/mL) containing 0.6nM Mouse Factor VII (Biotechne), 125nM Human Factor X (Prolytix), and 150µM Factor Xa chromogenic substrate (Sigma-Aldrich) was added to each well. Absorbance at 405nm was read every 3 minutes on a Biotek Synergy HTX microplate reader set to 37°C with a shaking step before each reading. Absorbance readings were blanked to the absorbance of the well at 0 minutes.

### Statistics

GraphPad Prism was used to calculate statistical significance. Statistical tests are indicated in the figure legends. *,**,***,**** corresponds to 0.01>P>0.05, 0.001>P> 0.01, 0.0001>P>0.001, P< 0.0001 respectively.

### Study approval

All procedures and use of animals was approved by the University of Washington IACUC.

### Data and material availability

Values for all figures can be found in the supplemental data values file. Reagents described in this manuscript are available from A.O. for research use under a materials transfer agreement from the University of Washington.

## AUTHOR CONTRIBUTIONS

Project conceptualization by C.H., A.G.S., and A.O.; Preliminary data generated by A.G.S. and C.H.; Experimental design by C.H. and A.O.; Experiments conducted and analyzed by C.H.; Technical advice and project guidance provided by M.H.; Cell lines generated by C.H. and A.G.S.; Manuscript written by C.H. and A.O.; Manuscript edits by C.H., A.O., M.H., A.G.S.

## ACKNOWLEDGMENTS

Special thanks to Joanne Estergreen, Department of Laboratory Medicine and Pathology, UW Medical Center, for help designing the plasma coagulation assay. Flow cytometry data was acquired through the University of Washington, Cell Analysis Facility Shared Resource Lab, with NIH award 1S10OD024979-01A1 funding for the Symphony A3. Microscopy was performed at the W. M. Keck Microscopy Center with training from Dr. Nathaniel Peters. Graphics created with BioRender.com. This work was funded by NIH grant U01CA289564 (to AO and MH). C.H. was supported by T32 GM007270.

