## Supplemental Figures for "Apoptotic cells promote circulating tumor cell survival and metastasis"

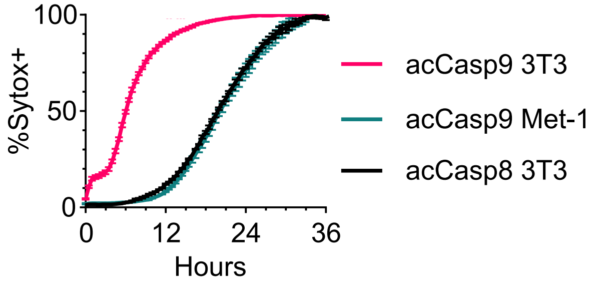


**Supplemental Figure 1. Cell Death Kinetics of activatable apoptosis systems.** acCasp9 3T3, acCasp9 Met-1 and acCasp8 3T3 were incubated with B/B and Sytox green dye. % Sytox+ cells were measured with Incucyte live cell imaging to measure membrane permeability.

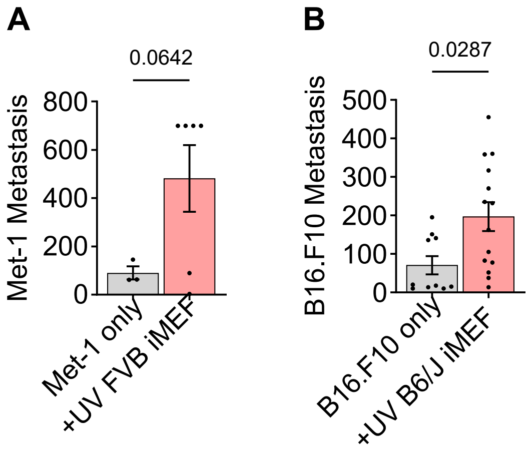


**Supplemental Figure 2. UV irradiated iMEFs promote metastasis.** MEFs were derived from FVB/NJ (A) or B6/J (B) mice and immortalized with SV40LT transduction (iMEF). iMEFs were UV irradiated and injected into mice at a 1:1 ratio with Met-1 (A) or B16.F10 (B). Surface lung nodules were quantified 14 days after I.V. injection. Lungs containing >700 surface metastasis were unable to be accurately quantified due to significant overlap of nodules, thus the maximum value of 700 was recorded in these cases. Dots are biological replicates, statistical testing is unpaired t-test.

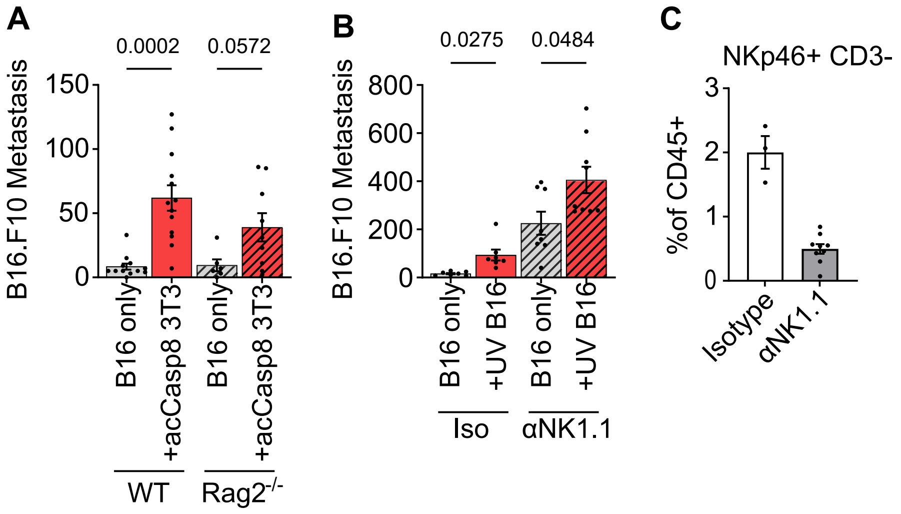


**Supplemental Figure 3. Effects of apoptotic cells on B16.F10 metastasis in Rag2$^{-/-}$ and NK cell depleted animals.** Apoptotic cells were injected at a 1:1 ratio with B16.F10 tumor cells. Surface lung nodules were quantified 14 days after I.V. injection (A-B). NK cell frequency of total CD45+ cells in the blood was measured 24 hours after I.V. injection of tumor cells to confirm NK cell depletion (C). Dots are biological replicates analyzed by ordinary one-way ANOVA with Tukey's multiple comparisons test.

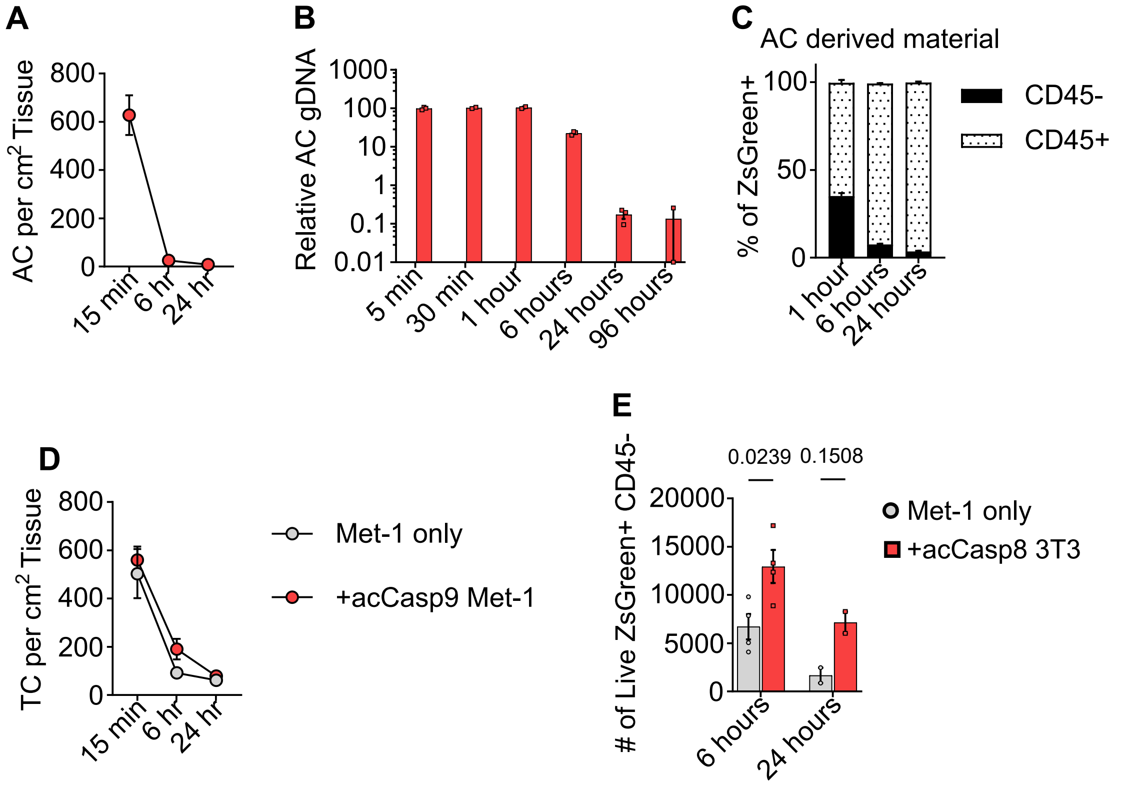


**Supplemental Figure 4. Quantification of Apoptotic cell and Tumor cell persistence in the lung.** Mice were injected with Met-1 cells and acCasp9 Met-1 (A,D) or acCasp8 3T3 (B,C,E) and lungs harvested at various timepoints for analysis by fluorescent microscopy (A,D), qPCR (B), or Flow Cytometry (C,E). Tumor cells expressed ZsGreen (A,D,E) and apoptotic cells expressed mCherry (A,D,E) or ZsGreen (B,C). Dots are biological replicates, statistical testing is unpaired t-test. AC=Apoptotic Cell, TC=Tumor Cell

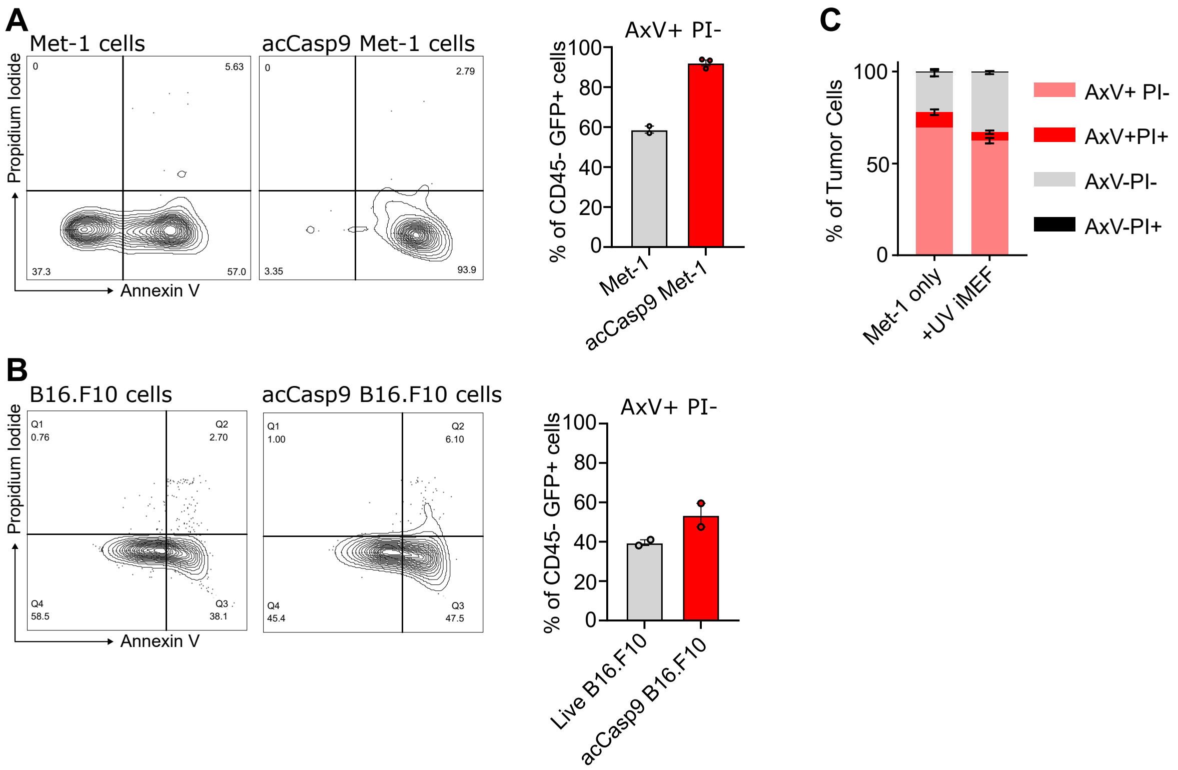


**Supplemental Figure 5. Evidence of phosphatidylserine exposure on tumor cells 1 hours after I.V. injection.** Viable Met-1 or B16.F10 cells were injected I.V. into mice and lungs were harvested and processed for flow cytometry. acCasp9 Met-1 or acCasp9 B16.F10 were activated with B/B for 15 minutes prior to I.V. injection and served as a positive control for PS externalization (A-B). Met-1 tumor cells were injected alone or at a 1:1 ratio with UV irradiated iMEFs(C). Lungs were processed for flow cytometry and stained with fluorescent Annexin V (AxV) and PI. Tumor cells expressed ZsGreen or GFP and were gated on as GFP+, CD45- population after excluding doublets and debris.

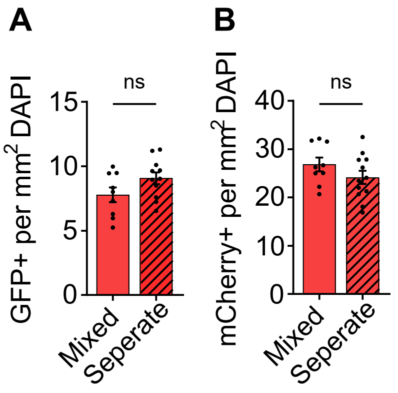


**Supplemental Figure 6. Fluorescent imaging quantification of tumor cell and apoptotic cell seeding.** Met-1 cells (mCherry+) and acCasp9 Met-1 (GFP+) were mixed up to one hour in advance (mixed) or injected in two sequential injections into opposite tail veins (separate). Lungs were harvested 1 hour after I.V. injection for fluorescent imaging. Error bars represent SEM, multiple slices of lung from n=3-4 biological replicates were analyzed with unpaired T-tests (A-B) or ordinary one-way ANOVA with Tukey's multiple comparisons test

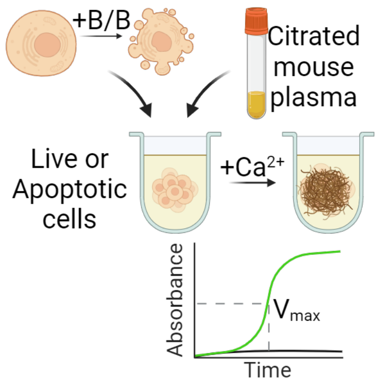


**Supplemental Figure 7. Diagram of plasma coagulation assay.** Live cells or apoptotic cells activated with B/B are added to citrated mouse plasma. $Ca^{2+}$ is added and the absorbance at 405nm is measured over time. The time where the maximum change in absorbance is observed is calculated and represents how fast a fibrin clot forms.

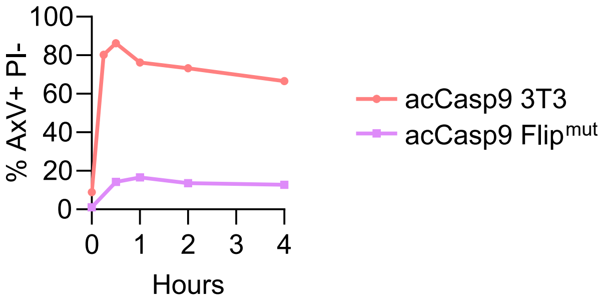


**Supplemental Figure 8. Kinetics of phosphatidylserine externalization after acCasp9 activation in parental or Flip$^{mut}$ cells.** Cells were activated with B/B and analyzed by flow cytometry at various timepoints, staining for PS exposure with Annexin V (AxV) and membrane integrity with propidium iodide (PI).

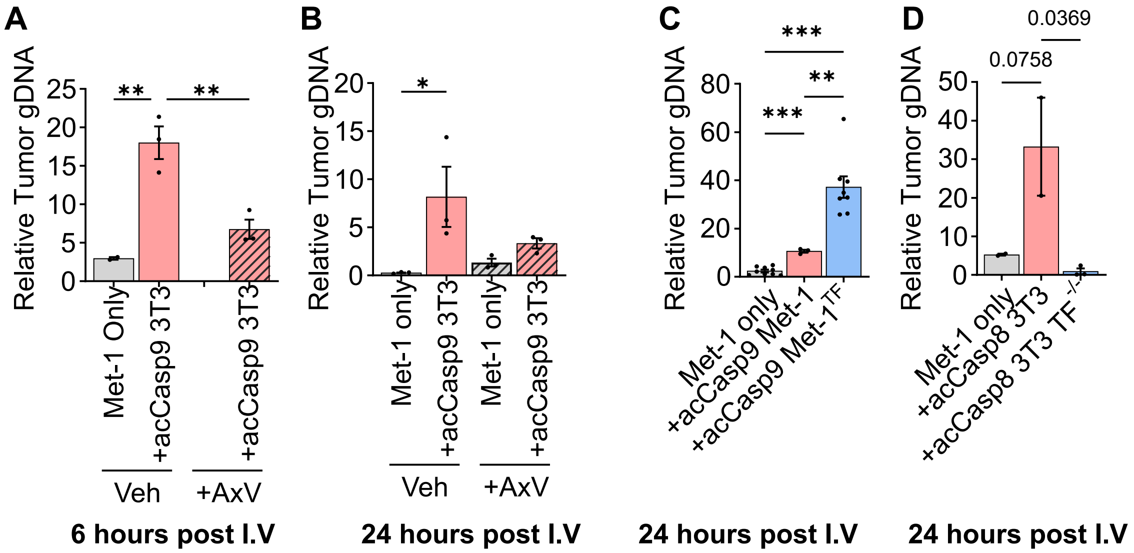


**Supplemental Figure 9. qPCR quantification of tumor cell persistence in lungs.** Mice were injected with Met-1 cells alone or with Met-1 cells and various types of apoptotic cells. Cells were resuspended in HBSS +$Ca^{2+}$ +$Mg^{2+}$ with or without Annexin V (AxV) (A,B) or resuspended in PBS (C,D). Lungs were harvested at either 6 hours (A) or 24 hours (B-D) and tumor cell gDNA was isolated and quantified by qPCR.

| Target | Fwd Primer | Rev Primer | FAM/NFQ-MGB Probe |
|---|---|---|---|
| **Ptger2** | TAC CTT CAG CTG TAC GCC AC | GCC AGG AGA ATG AGG TGG TC | /56-FAM/CC TGC TGC T/ZEN/T ATC GTG GCT G/3IABkFQ/ |
| **ZsGreen** | GTA CCA CGA GTC CAA GTT CTA C | CAC GTC GCC CTT CAA GAT | /56-FAM/CC CGT GAT G/ZEN/A AGA AGA TGA CCG ACA A/3IABkFQ/ |
| **PyMT** | CGA AAT CCT TGT GTT GCT GA | GCT GGT CTT GGT CGC TTT C | /56-FAM/CC GAT GAC A/ZEN/G CAT ATC CCC /3IABkFQ/ |
| **mCherry** | GAC TAC TTG AAG CTG TCC TTC C | CGC AGC TTC ACC TTG TAG AT | /56-FAM/TT CAA GTG G/ZEN/G AGC GCG TGA TGA A/3IABkFQ/ |
| **GFP** | GAA CCG CAT CGA GCT GAA | TGC TTG TCG GCC ATG ATA TAG | /56-FAM/AT CGA CTT C/ZEN/A AGG AGG ACG GCA AC/3IABkFQ/ |
| **101a** | GAG GAG ACT GTA CGC AAG ATG | TGG CGC TGC TGT TTG ATA | /56-FAM/AT CAC CA C C/ZEN/T TCA CCT CGT TGC C/3IABkFQ/ |

**Supplemental Table 1. qPCR Primer/Probe Sets**
